# Mitigating ecosystem impacts of bottom trawl fisheries for North Sea sole *Solea solea* by replacing mechanical by electrical stimulation

**DOI:** 10.1101/2020.01.21.913731

**Authors:** A.D. Rijnsdorp, J. Depestele, O.R. Eigaard, N.T. Hintzen, A. Ivanovic, P. Molenaar, F. O’Neill, H. Polet, J.J. Poos, T. van Kooten

## Abstract

Ecosystem effects of bottom trawl fisheries are a major concern. We analysed whether the replacement of mechanical stimulation by electrical stimulation may reduce the adverse impacts on the benthic ecosystem in the beam trawl fishery for sole. Although the use of electricity is not allowed to catch fish in European Union waters, a number of beam trawlers got derogation and switched to pulse trawling to explore the potential to reduce impacts. We extended a recently developed assessment framework and showed that the switch to pulse trawling substantially reduced benthic impacts when exploiting the Total Allowable Catch of sole in the North Sea. We applied the framework to Dutch beam trawl logbook data from 2009 to 2017 and estimated that the trawling footprint decreased by 23%; the precautionary impact indicator of the benthic community decreased by 39%; the impact on median longevity decreased by 20%; the impact on benthic biomass decreased by 61%; the amount of sediment mobilised decreased by 39%. The decrease is due to the replacement of tickler chains by electrode arrays, a lower towing speed and higher catch efficiency for sole. The effort and benthic impact of the beam trawl fishery targeting plaice *Pleuronectes platessa* in the central North Sea increased with the recovery of the plaice stock. This study illustrates the usefulness of a standardized methodological framework to assess the differences in time trends and trawling impact between gears.

## Introduction

Ecosystem effects of bottom trawl fisheries are a major concern [1–3]. Bottom trawling takes place over large parts of the continental shelves and is responsible for about 25% of the wild marine landings [4, 5]. Bottom trawling generally requires heavy fishing gears and powerful engines with a high fuel consumption and CO2 emission [6]. Bottom trawling homogenises sea floor texture, disturbs the sorting of sediment generated by natural or biological processes [7–9], mobilises fine sediments into the water phase [10, 11], and may cause sediment systems to become unstable [12]. Further, bottom trawls impact benthic communities by damaging habitats and by imposing direct mortality to benthic animals [13–15]. All these impacts also affect bio-geochemical processes in the sea floor – water interface and food webs [16] [11] [17].

Beam trawls used to target flatfish species, in particular sole (*Solea solea*), are considered to be among the fishing gears with the largest ecological impact on the benthic ecosystem[14]. The tickler chains dragged over the sea floor to chase sole into the net, penetrate the sediment and disturb the top layer of the sea bed down to a depth of 4 - 8 cm [18–20]. The relatively small cod-end mesh size required to retain the slender soles, results in large bycatches of undersized plaice and other fish species [21–23]. Since the introduction of the beam trawl in the 1960s, fishers have invested in larger vessels to increase gear size, towing speed, and the number of tickler chains [24]. This increase in fishing capacity fuelled concern about the environmental impacts of this fishery [25].

Already in the 1970s, research started to investigate the possibility to replace mechanical stimulation using tickler chains by electrical stimulation in the beam trawl fishery for flatfish [26]. It was shown that electrical stimulation can be deployed to immobilise fish, preventing them to escape from the approaching gear. After a successful year-round trial in 2004 with a commercial prototype [27], many vessels switched to pulse trawling for sole between 2009 and 2015. The successful introduction was related to the improved selectivity and catch efficiency for the main target species sole [28, 29] and a reduction in fuel consumption due to reduced towing speed [6, 28]. Because EU-legislation does not allow the use of electricity to catch fish, pulse trawlers operate under a (temporary) derogation [30]. To support decision making on the question whether pulse trawling can be accepted as a legal fishing method [31], information is required on the ecosystem impacts of both the traditional beam trawl gear and the innovative pulse trawl.

In this paper we study whether a transition from traditional beam trawling to pulse trawling can reduce the impact of trawling on the benthic ecosystem, focussing on the consequences of mechanical disturbance. We apply a recently developed impact assessment framework [14, 32–34] to estimate fishing footprints (areal extent) and benthic impact indicators, based on distribution maps of the fishery and dimensions of the fishing gears [5, 35] and the sensitivity of the benthic community[36, 37]. In addition to indicators for precautionary impact (L1); median longevity of community (L2); and community biomass (PD), we estimate the amount of fine sediments mobilised in the turbulent wake of the fishing gears.

## Material and methods

### Beam trawl fleet

The Dutch beam trawl fleet use two outriggers to deploy a beam trawl from each side of the vessel when trawling for flatfish in the North Sea. The width of a beam trawl is restricted to 12 m for vessels with engine power > 221kW and 4.5m for vessels with a maximum engine power of 221kW when fishing in coastal waters. The minimum mesh size allowed is 80mm in the sole fishing area (SFA) in the south and 100mm in the plaice fishing area in the North. The border between SFA and North is determined by a demarcation line running from west to east at 55°N shifting to 56°N east of 5°E.

The horizontal net opening of a beam trawl is fixed by a beam that rest on two shoes (Fig 1a). Since 2008, most vessels replaced the beam and shoes by a hydrodynamic wing (Fig 1b). The use of the innovative SumWing reduced the fuel consumption by 16% due to the improved streamline and the reduced bottom contact[6].

**Fig 1.**
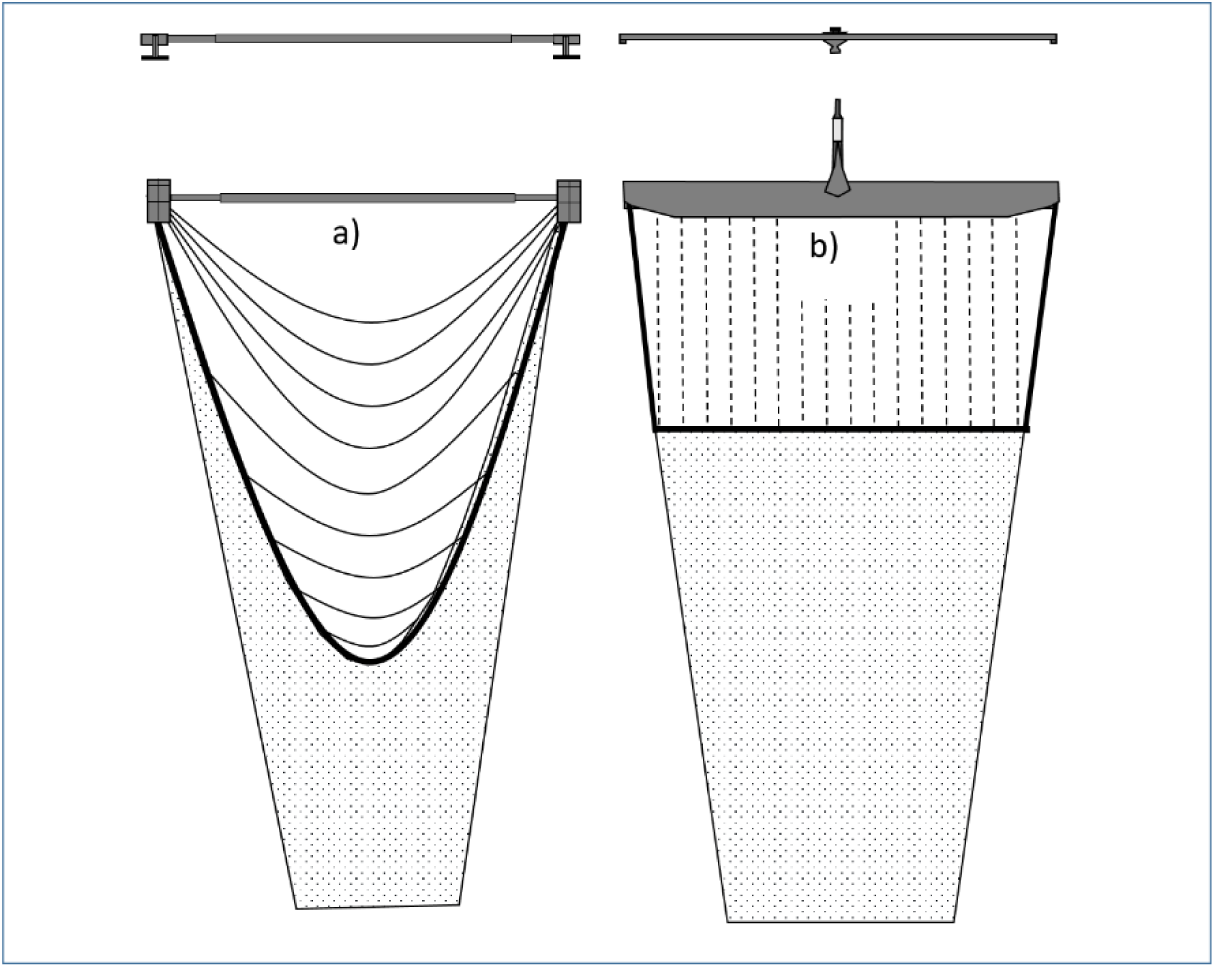
Schematic drawing of the tickler chain beam trawl (left) and the pulse trawl (right). For each gear the frontal (top) and bottom (middle) view of the beam or wing is shown as well the rigging of the tickler chains and ground rope, or electrode arrays and ground rope. Bottom contacting gear components are the shoes, tickler chains and ground rope of the tickler chain beam trawl, and the nose, ground rope and electrode arrays of the pulse trawl.

To chase the flatfish up from the sea bed, the conventional tickler chain beam trawl deploys a row of transverse tickler chains that are attached to the shoes and ground rope (Fig 1a) [38]. In pulse trawls the tickler chains are replaced by a rectangular array of electrodes that is fitted between the beam/wing and ground rope and run parallel to the towing direction.(Fig 1b)[39, 40]. In order to fit this rectangular array of electrodes, a horizontal ground rope is created by deploying a number of tension relief cords between the beam/wing and ground rope [19, 40]. In contrast to the electrode arrays, the tension relief cords do not have contact with the sea bed (video footage; Polet, pers comm).

### Catch and effort data

Two data sets of catch and effort (landings by species, hours at sea, ICES rectangle, gear, mesh size, vessel ID, landing date) by fishing trip of Dutch flagged vessels were extracted from the VISSTAT data base of mandatory logbooks for the period 2009 – 2017. Set 1 comprises all fishing trips of vessels that have reported to use a beam trawl gear (TBB) targeting flatfish. Set 2 comprises the data from a subset of vessels referred to as the set of pulse license holders (PLH). A maximum of 74 pulse trawlers have been active at the same time. Because a separate code for pulse trawl fishing trips (PLH_pulse) was not available for the full study period, pulse fishing trips were identified based on the reported mesh size (70-99mm), mean towing speed during fishing, and the start date of the pulse license (data LNV) [29].

### VMS data

Vessel speed and vessel positions of all beam trawl vessels were available from the Vessel Monitoring by Satellite (VMS) program. Vessel speeds typically show a three modal frequency distribution which allows us to distinguish the fishing positions from the positions during steaming or during floating [41, 42].

### Habitat variables

Habitat variables (%gravel, %mud) were obtained for 1×1 minute grid cells from [43]. Tidal bed shear stress (N.m-2) was obtained from a hydrodynamic model by John Aldridge (CEFAS) as used in [44] and [45].

### Trawling impact indicators

#### Footprint and trawling intensity

VMS fishing positions were interpolated to estimate the swept area by 1×1 minute grid cell longitude and latitude [46] and the trawling intensity is expressed by the swept area ratio. The grid cell resolution corresponds to approx. 1.9 km2 at 56° N with cell size gradually increasing/decreasing the further south/north it is located. At this resolution bottom trawling can be considered to be randomly distributed within a grid cell on an annual basis [4, 47, 48] and to become uniform at longer time scales [49].

Following [5], the trawling footprint was estimated as (i) the total surface area (km^2^) trawled at least once a year under the assumption of a uniform distribution of trawling activities within a grid cell, and (ii) the proportion of grid cells with any trawling activity irrespective of the trawling intensity. The latter metric includes the untrawled part of fished grid cells.

#### Sediment mobilization

Sediment mobilisation *m* is calculated from hydrodynamic drag H_d_ caused by the fishing gear and the silt fraction sf of the sediment [50, 51].

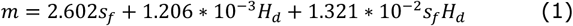

The hydrodynamic drag of the various types of beam trawls and pulse trawls is estimated from a quantitative inventory of the gear types and corresponding dimensions of the major gear elements [40] (Table 1).

**Table 1.** Estimated hydrodynamic drag (H_d_ in 10^3^ N.m^-1^) of different types of beam trawls used in the flatfish fisheries in the North Sea (from[40]).

| Type of beam trawl | Euro cutters | Large vessels |
| --- | --- | --- |
| Tickler chain beam trawl | 2.8 | 6.2 |
| Pulse trawl | 2.9 | 3.8 |

#### Impact

Three methods were used to assess the impact of bottom trawling on the benthic ecosystem (reviewed in [52]). All three methods build on the assumption that the sensitivity of the benthos for bottom trawling is related to the longevity composition of the benthic community which can be described by the cumulative biomass (B) as a function of ln(longevity) (L), habitat (H) and trawling (T) [36, 37].

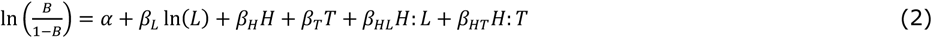

#### Precautionary approach (L1)

L1 estimates the proportion of the biomass of the benthic community that is potentially impacted by trawling [33]. It assumes that benthic taxa with a longevity of more than the average interval between two successive trawling events will be potentially affected by bottom trawling. Hence the impact can be estimated as the proportion of biomass of those taxa with a longevity exceeding the reciprocal trawling intensity (L = 1/T), which was derived from equation [1] as:

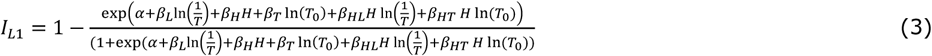

Because the impact is estimated relative to the untrawled community, a value of T0 = 0.01 was included to avoid taking the log of zero.

#### Statistical-impact approach (L2)

Trawling shifts the community composition towards shorter lived taxa. The median longevity of the community *M*_*T*_in response to trawling is based on the statistical relationships between trawling intensity and longevity as found in [37].

By re-arranging equation (2), *M*_*T*_is given by:

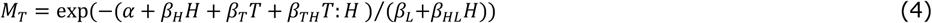

L2 estimates the relative change in median longevity in response to trawling by:

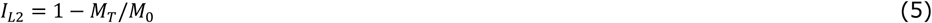

where *M*_*T*_ is the median longevity at trawling intensity T and *M*_0_ is the median longevity of the untrawled community.

#### Population dynamic approach (PD)

The population dynamic approach estimates the impact of bottom trawling (I) in terms of the reduction in the benthic biomass (B) relative to the carrying capacity (K) of the habitat [32, 36]

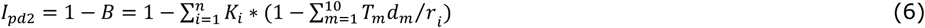

Where *r*_*i*_ is the recovery rate and *K*_*i*_ is the biomass proportion of longevity class i in the total community, and *T_m_* is the trawling intensity and *d_m_* is the depletion rate of métier *m*.

#### Parameterisation

The parameters of the longevity composition in relation to habitat variables and trawling intensity (eq 1) are based on [37]. The depletion rate of the tickler chain beam trawl (dT=0.14) is based on the results of the meta-analysis of[14]. Given the observed linear relationship between depletion rate and penetration depth across gears [14]and the 50% reduction in penetration depth of the pulse trawl relative to the tickler chain beam trawl[19], the depletion rate of the pulse trawl was estimated as d_P_=0.5*d_T_. The recovery rate was set at *r* = 5.31*longevity^-1^[36]. The number of longevity classes used in the calculations was set at n=10.000 with a maximum longevity of 100 years.

## Results

### Towing speed

Pulse trawls are towed at a 23% and 13% lower speed than tickler chain beam trawls in large and small vessels, respectively (Table 2, Fig 2),

**Fig 2.**
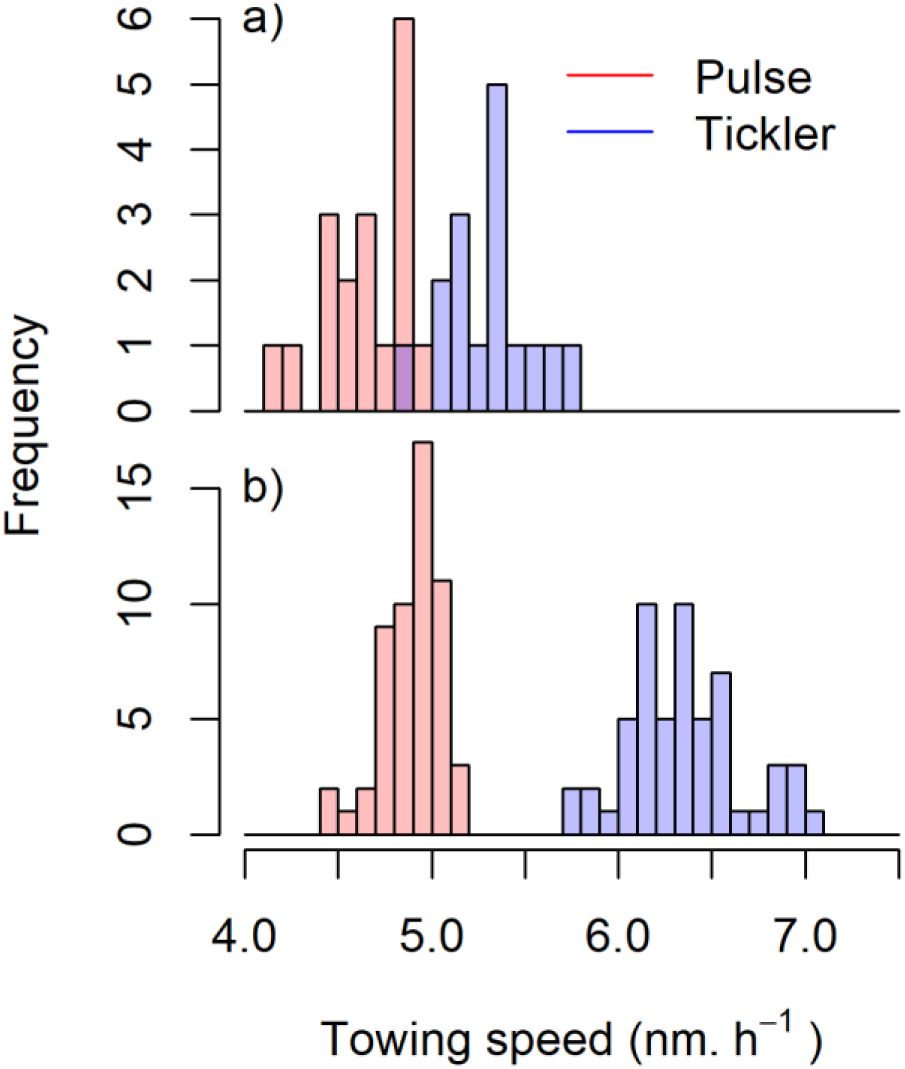
Histogram of the mean towing speed of small (< =221 kW) and large vessels (>221 kW) using a tickler chain or pulse trawl. Towing speeds are estimated from the VMS recorded speed of PLH.

**Table 2.**
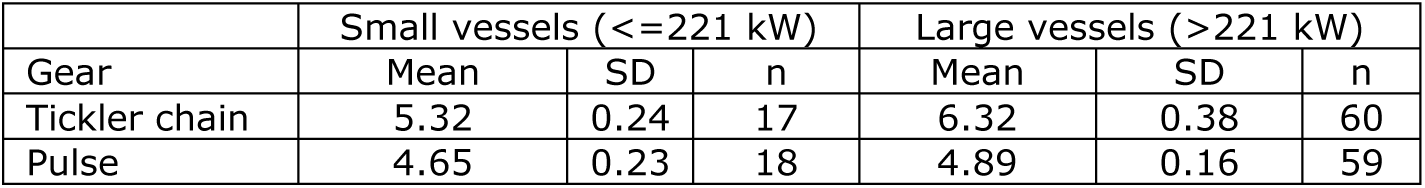
Mean towing speed of pulse licence holders when fishing with the traditional tickler chain beam trawl or pulse trawl.

### Effort and landings

Trends in fishing hours and landings of the total beam trawl fleet (thin lines) and subset of PLH (thick lines) are shown in Fig 3 for the total North Sea (full lines) and for the sole fishing area (SFA) south of the demarcation line (hatched lines). The fishing hours of the Dutch beam trawl fleet decreased from 470 thousand in 2009 to 347 thousand in 2014 and then increased to 394 thousand in 2017. Most beam trawling occurs in SFA south of the demarcation line. Pulse trawl license holders maintained their fishing effort targeting sole in SFA at around 310 thousand hours during the transition, but increased their effort targeting plaice north of SFA from 10 thousand hours in 2009 to 40 thousand hours in 2017. The contribution of PLH to the fishing hours of the Dutch beam trawl fleet increased from 66% in 2009 to 86% in 2017.

**Fig 3.**
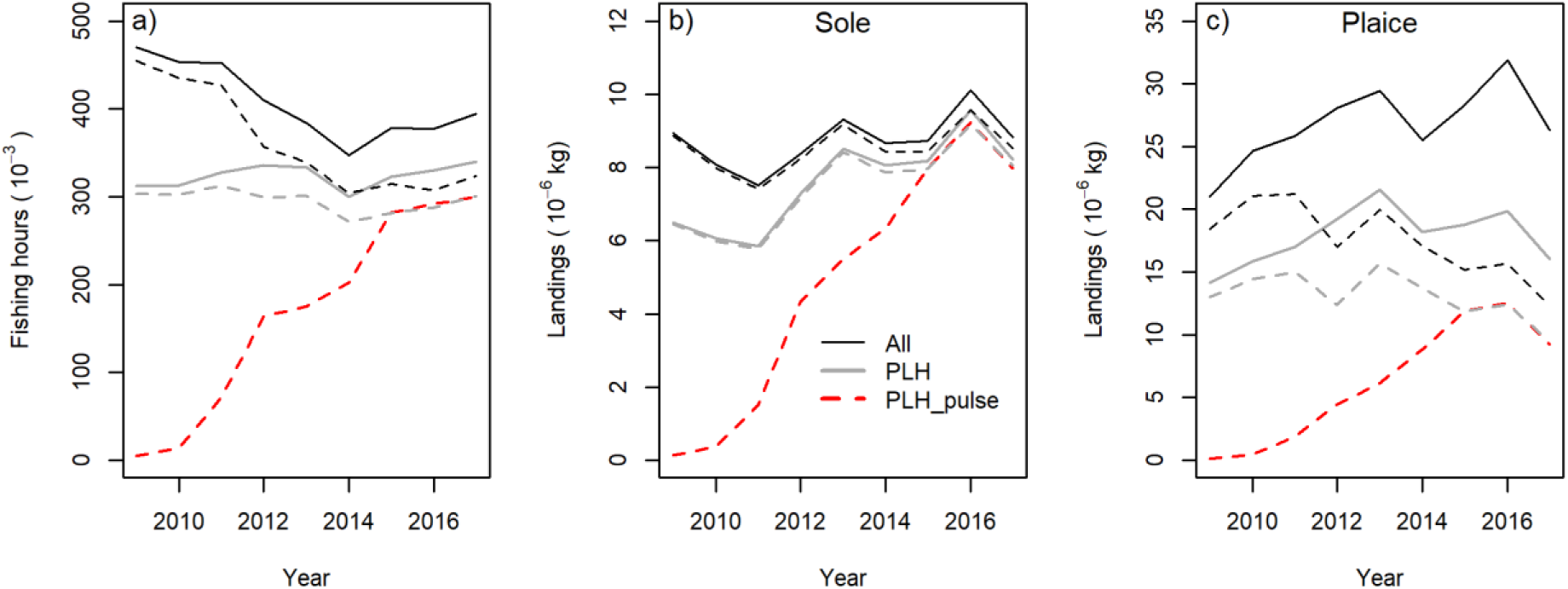
Changes in fishing effort (fishing hours) and landings of sole and plaice of the Dutch beam trawl fleet (ALL) and the subset of pulse license holders fishing with a tickler chain or pulse trawl (PLH) and fishing with a pulse trawl (PLH-pulse). Plaice landings also include landings with the twin otter trawl. Full lines refer to the total fishing area. Hatched lines refer to the sole fishing area (SFA) with 80mm mesh size south of the demarcation line.

Annual sole landings of the Dutch fleet varied between 8 and 10 thousand tons. The contribution of the PLH to the Dutch sole landings increased from 73% in 2009 to 93% in 2017 (Fig 3b). The plaice landings increased during the study period from 20 to 25-30 thousand ton (Fig 3c). The proportion of plaice landed by the PLH slightly decreased from 67% in 2009 to 61% in 2017 of which two-third was landed by the pulse trawl and one-third by the tickler chain trawl. The proportion of plaice landed from SFA decreased from close to 100% in 2009 to about 50% in 2017.

### Spatial distribution

Fig 4 compares the spatial distribution of the trawling (swept area ratio) of the Dutch beam trawl fleet before (2009-2010) and after (2016-2017) the transition to pulse trawling. In 2009-2010 the Dutch beam trawl fleet mainly fished in the SFA. After the transition to the pulse trawl, the fleet continued fishing for sole in the SFA although changes in relative fishing intensity occurred. Within the SFA, trawling intensity was more or less stable south of the 53°N (IVc), except for a slight increase within the 12 nm zone of the Belgium coast, off the Thames estuary and parts of the Norfolk banks, and was reduced on the fishing grounds located between 53°N and 55°N. In the area north of SFA the beam trawl fleet increased its fishing activities targeting plaice with a 100 mm cod-end.

**Fig 4.**
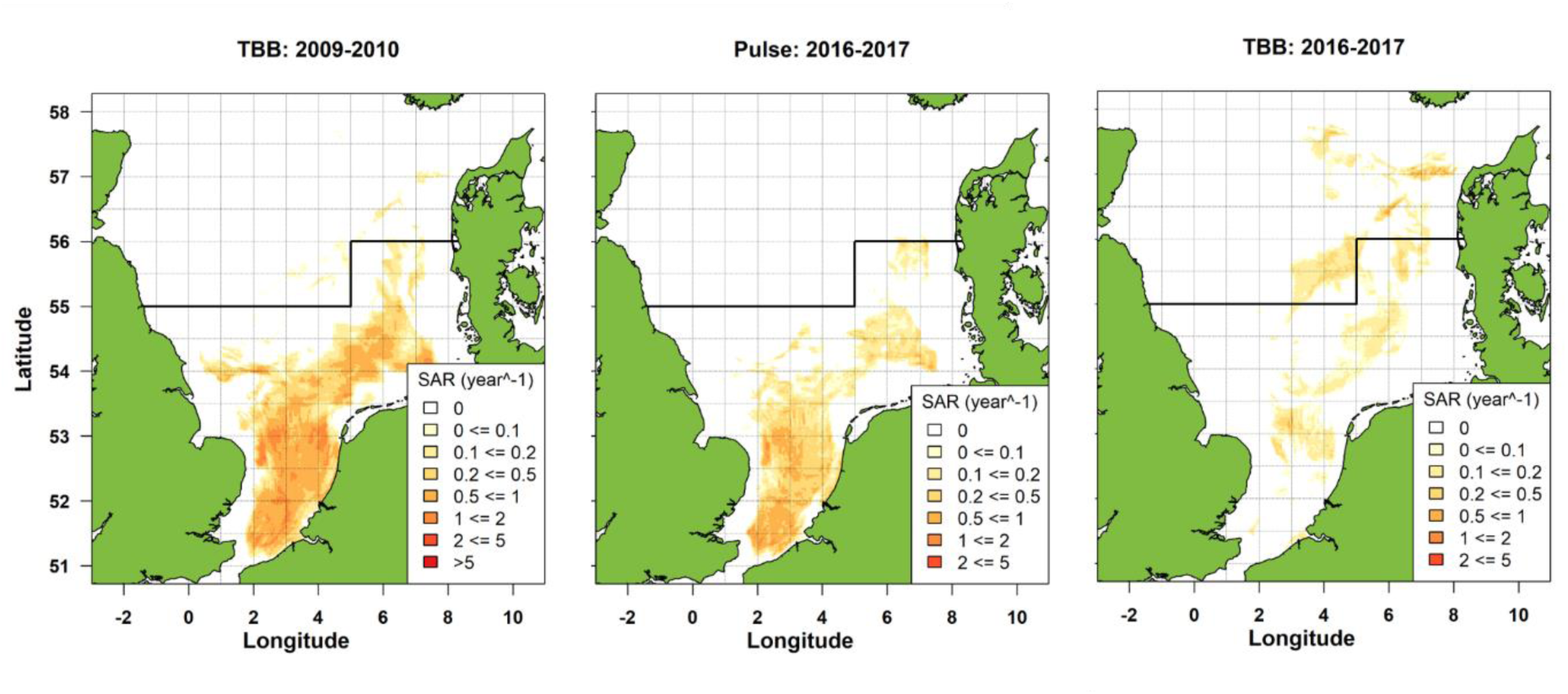
Spatial distribution of trawling intensity (annual swept area ratio SAR) of the total Dutch beam trawl fleet before (TBB 2009-2010) and after (Pulse 2016-2017 and TBB 2016-2017) the transition to pulse fishing. Pulse fishing is restricted to the sole fishing area (SFA) south of the demarcation line at 55oN and 56oN (minimum cod-end mesh size of 80 mm). North of the demarcation line a minimum mesh size of 100 mm is required and the beam trawling target plaice.

### Trawling footprint and habitat association

The area swept by the beam trawl fleet (fishing hours*gear-width*towing speed) decreased by about 33% between 2009 and 2014 and remained stable since then (Fig 5a). The area swept by the PLH showed a similar pattern but with a smaller decrease of about 21%. The decrease in swept area was particularly strong in SFA, 42% for the total fleet and 28% for the PLH. The decrease in swept area is due to both a decrease in fishing hours and towing speed.

**Fig 5.**
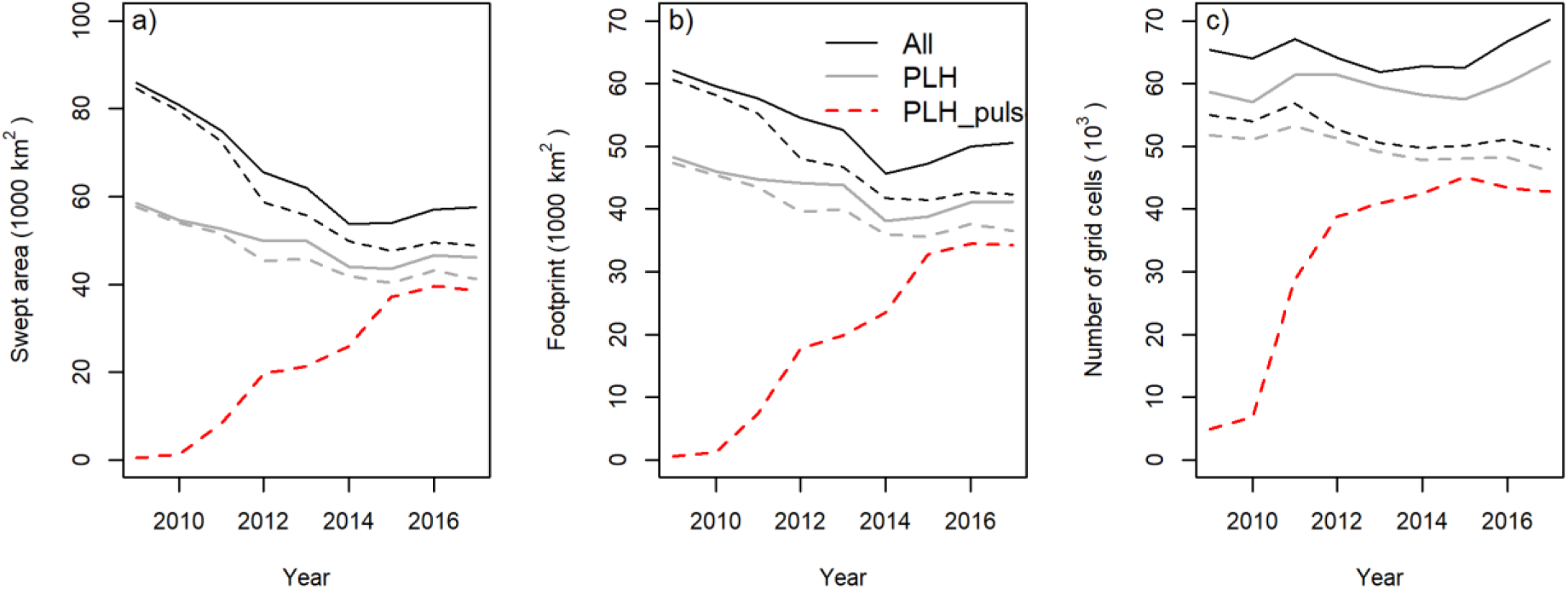
Changes in the area swept, the surface of the sea floor which is trawled at least once per year (footprint) and the number of 1×1 minute grid cells with trawling activities recorded for the total Dutch beam trawl fleet (ALL) and for the subset of pulse license holders fishing with a tickler chain trawl or a pulse trawl (PLH) or with a pulse trawl (PLH-pulse). Full lines refer to the total fishing area. Hatched lines refer to the sole fishing area (SFA) with 80mm mesh size south of the demarcation line.

The annual footprint of the beam trawl fisheries, defined as the surface area of the sea floor that is trawled at least once in a year, decreased during the transition by 19% from about 62 thousand km^2^ in 2009 to 50 thousand km^2^ in 2017 (Fig 5b). The decrease was less than the decrease in swept area. The footprint of the PLH, including pulse and tickler chain trawling, decreased by 15% from 48 thousand km2 in 2009 to 41 thousand km^2^ in 2017. After the transition, the footprint of the pulse trawl varied around 34 thousand km^2^. The number of 1×1 minute grid cells with trawling activities varied without a clear trend (Fig 5c), although the number of grid cells in 2017 was 7% higher in the total fishing area and 10% lower in SFA than in 2009. The number of grid cells with pulse trawl activities reached a stable level in 2012 when the swept area only reached about half of its final level in 2015 and later years (Fig 5a).

The habitat association of the beam trawl fleet is presented in Table 3. The Dutch beam trawl fleet deploys more than 80% of its fishing effort on sandy sediments which comprise only 60% of sea floor habitats in the SFA. Coarse and mixed sediments are fished less than their proportional occurrence, while mud is trawled in proportion to its occurrence. Pulse trawling occurs slightly more in coarse habitats and less in mud.

**Table 3.**
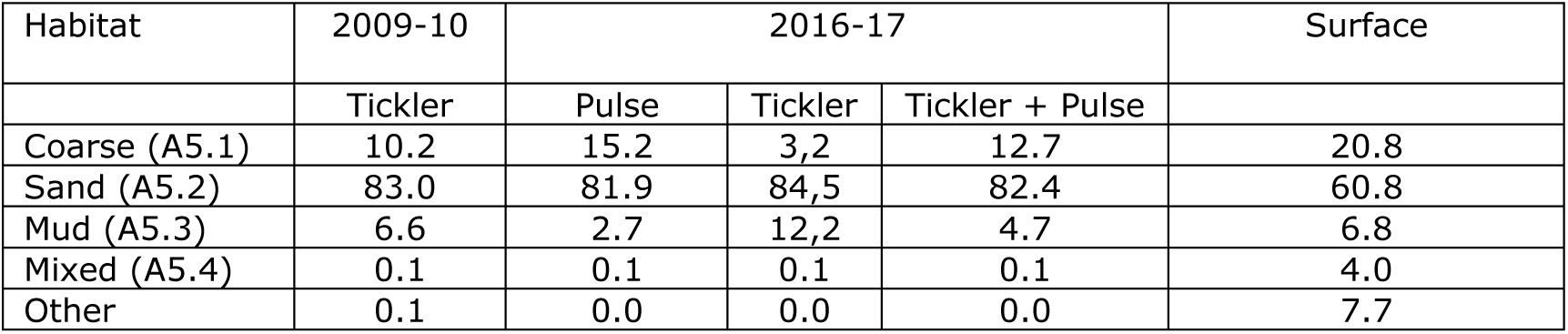
Percentage fishing effort (swept area) of the Dutch beam trawl fleet and percentage surface area by Eunis habitat in the sole fishing area (SFA) south of the demarcation line at 55oN and west of 5oE and 56oN east of 5oE.

| Habitat | 2009-10 | 2016-17 |  |  | Surface |
| --- | --- | --- | --- | --- | --- |
|  | Tickler | Pulse | Tickler | Tickler + Pulse |  |
| Coarse (A5.1) | 10.2 | 15.2 | 3,2 | 12.7 | 20.8 |
| Sand (A5.2) | 83.0 | 81.9 | 84,5 | 82.4 | 60.8 |
| Mud (A5.3) | 6.6 | 2.7 | 12,2 | 4.7 | 6.8 |
| Mixed (A5.4) | 0.1 | 0.1 | 0.1 | 0.1 | 4.0 |
| Other | 0.1 | 0.0 | 0.0 | 0.0 | 7.7 |

### Impact

The changes in benthic impacts are shown in Fig 6 for the total Dutch fleet and the subset of PLH. Benthic impact in the SFA (hatched lines) is substantially higher than in the total fishing area (black lines) because most fishing occurred in the southern area. During the transition, impact decreased for both groups. The impact of the pulse trawling fishing (PLH_pulse = red line) increased but never reached the impact level of the beam trawl activities of the PLH prior to the transition (PLH = grey line).

**Fig 6.**
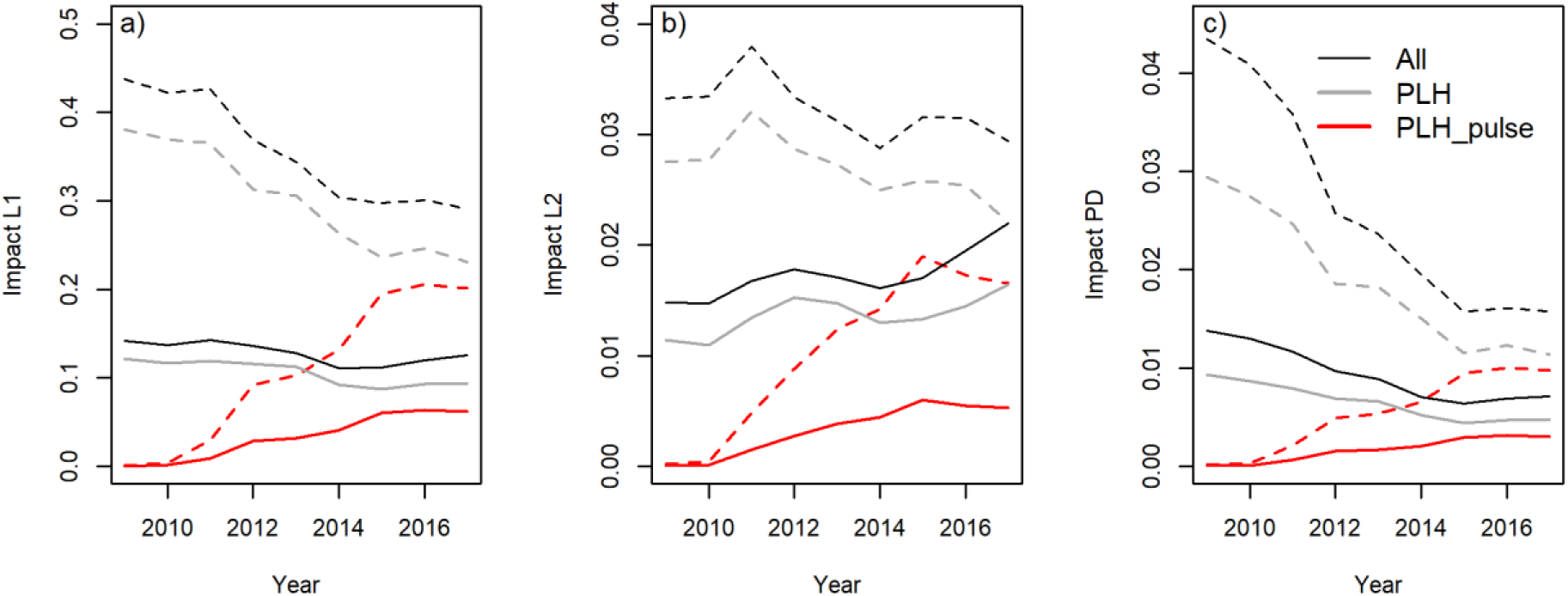
Time trends in the impact indicators of the total Dutch beam trawl fleet (ALL) and for the subset of pulse license holders fishing with a tickler chain trawl or pulse trawl (PLH) or with a pulse trawl (PLH-pulse). Full lines refer to the total fishing area. Hatched lines refer to the sole fishing area (SFA) with 80mm mesh size south of the demarcation line.

The L1 indicator, which estimates the proportion of the benthos with a life span exceeding the time interval between successive trawling events, decreased by 12% (SFA = 34%) for the total Dutch fleet and by 23% (SFA = 39%) for the PLH (Table 4).

**Table 4.** Ratio of the impact of the total Dutch beam trawl fleet and the subset of pulse license holders (PLH) in 2009 before and in 2017 after the transition to pulse trawling (I2017/I2009) in the total fishing area and the sole fishing area (SFA) south of the demarcation line. Values >1 indicate and increase in impact by pulse trawling.

| Indicator | Total fleet |  | Pulse license holders |  |
| --- | --- | --- | --- | --- |
|  | Total fishing area | Sole fishing area (SFA) | Total fishing area | Sole fishing area (SFA) |
| Swept area | 0.67 | 0.58 | 0.79 | 0.72 |
| Footprint | 0.81 | 0.70 | 0.85 | 0.77 |
| Number grid cells | 1.07 | 0.90 | 1.08 | 0.89 |
| Impact L1 | 0.88 | 0.66 | 0.77 | 0.61 |
| Impact L2 | 1.49 | 0.89 | 1.44 | 0.80 |
| Impact PD | 0.52 | 0.36 | 0.51 | 0.39 |
| Sediment mobilization | 0.41 | 0.34 | 0.67 | 0.61 |

The L2 indicator, which estimates the decrease in median longevity of the benthic community due to trawling, shows a gradual 11% decrease in the SFA for the total fleet as well as a 20% decrease for the PLH 20% (Table 4). When estimated for the total fishing area, however, an increase in impact is estimated of 49% and 44% for the PLH and the total fleet, respectively. The increase in the beam trawling with tickler chain trawls targeting plaice north of SFA where natural disturbance is low overrides the impact reduction due to the transition to pulse trawling in the SFA.

The Biomass indicator, which measures the decrease in equilibrium benthic biomass due to the trawling intensity, shows a clear decreasing trend in the SFA to a level in 2017, which is about 60% lower than in 2009 for both the total beam trawl fleet and the PLH. For the total fishing area, the decrease in impact is estimated at about 50% (Table 4).

### Sediment mobilization

The estimated amount of sediment that is mobilized in the wake of the beam trawls is estimated at 20.10^14^ kg.year^-1^ and decreased during the transition period (Fig 7). For the total fleet the amount is 59% (SFA = 66%) lower in 2017 than in 2009 (Table 4). For the PLH the decrease is 33% (SFA = 39%). After the transition in 2017, pulse trawl and tickler chain activities have an about equal share of the total amount of 8.10^14^ kg.year^-1^ sediments mobilized.

**Fig 7.**
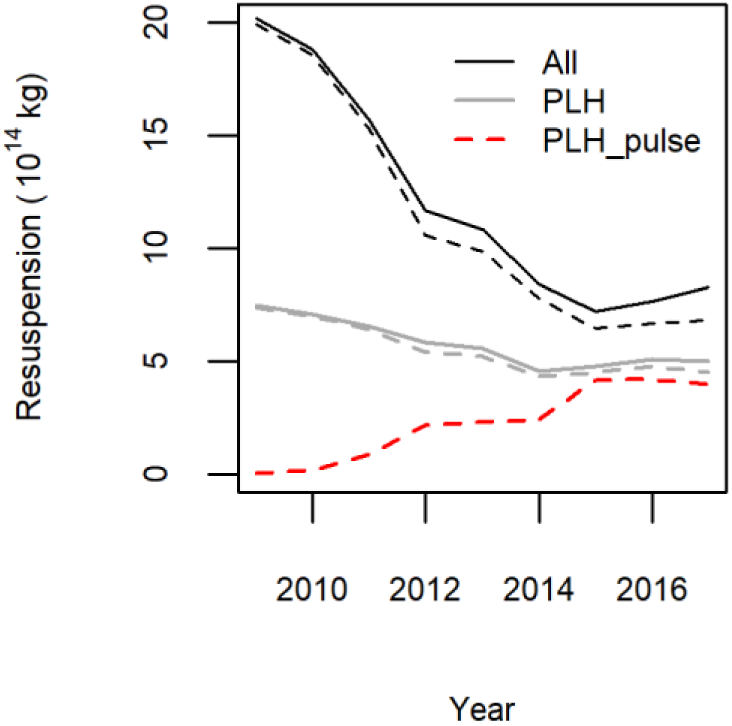
Time trends in the amount of sediments mobilised by the Dutch beam trawl fisheries (ALL: thick black), the subset of pulse license holders fishing with a tickler chain trawl or pulse trawl (PLH: thin black) or fishing with a pulse trawl (PLH-pulse: red). Full lines refer to the total fishing area. Hatched lines refer to the sole fishing area (SFA) with 80mm mesh size south of the demarcation line.

## Discussion

Pulse trawlers have been operating with a (temporary) exemption from the European Union ban on fishing and were allowed to do so as an exploration whether pulse trawling could reduce the ecological impacts of the traditional beam trawl fishery. To accommodate the interest of the Dutch fishing industry the Dutch government successfully negotiated an increase in licenses with the condition that the vessels would participate in research to assess the sustainability of the fishery [30]. Of the 84 available licenses, 76 were used for vessels in the sole fishery. Before the shift to pulse trawling these vessels accounted for about 73% of the sole landings and after the transition this share increased to about 95%. Hence, our study represents a full-scale experiment on the transition from tickler chain beam trawling to pulse trawling for sole, which not only allows an analysis of the transition consequences at the level of the individual vessel but also at the level of the fleet.

The transition to pulse trawling reduced the impact of the pulse license holders (PLH) on the benthic ecosystem between 20% and 61% in the SFA depending on metric (Table 4). The impact reduction of the PLH is a minimum estimate because the PLH increased their fishing rights for sole which to compensate for the increased catch efficiency (Poos in [53]) replacing fishing effort of other beam trawl vessels. On the other hand the impact reduction estimated for the total Dutch fleet is an overestimate because the beam trawl effort decreased due to vessels switching to fuel saving fishing gears such as twin otter trawl or flyshoot, or vessels leaving the fishery.

The reduction in impact is mainly due to two factors. First, electric stimulation allowed fishers to reduce towing speed and at the same time increase catch efficiency for sole, their main target species, but not for plaice (Poos in [53]). The increased catch efficiency for sole is likely due to its cramp response to electrical pulses, where it bends into a U-shape that can easily pass over the ground rope into the net [27, 54]. When exposed to a pulse stimulus, plaice also cramps, but does not bend noticeably and may pass underneath the ground rope.

Second, the replacement of transverse rows of tickler chains with longitudinal arrays of electrodes reduces the contact area of the trawl with the sea floor. In contrast to the tickler chains, that disturb the sea floor over the full width of the trawl, the contact area of a pulse trawl is restricted to the nose of the wing and the electrode arrays that run parallel to the towing direction [40]. In addition, the sediment penetration depth of the pulse trawl components is less [20]. In a comparative trawling experiment in fine sand it was shown that a tickler chain trawl disturbed the sea bed to a median depth of 4.1 cm, more than twice the median disturbance depth (1.8 cm) of a pulse trawl[19].

The reduced bottom contact of the pulse trawl implies reduced catch efficiency for benthos. Van Marlen et al (2014) showed that the amount of benthos caught in pulse trawls was 20% lower than in a tickler chain beam trawl fishing on the same grounds. In addition we expect that the reduced bottom contact and the lower towing speed will reduce the mortality caused by the physical contact with the gear [55]. Only two experimental studies have compared the impact of pulse trawls and tickler chain beam trawls, with equivocal results. In a BACI experiment in the Frisian Front area the depletion of benthos averaged over all species was lower for pulse trawling (25%) than for tickler chain trawling (44%) although the difference was not statistically significant[56]. In a study in coarser sediment in coastal water, where the benthic community mainly consisted of species that can be considered to be resistant to bottom trawling, no significant effect of beam trawling with either gear type could be detected [57].

The equivocal results of the two experiments are not surprising because it is notoriously difficult to quantify the trawling induced mortality in field experiments due to the generally large variance in the data [13]. A meta-analysis of the available studies, however, showed that the depletion rate is related to the penetration depth of the gear [13, 14]. The measured reduction in penetration depth of the pulse trawl of about 50% [19] and proportional reduction in depletion rate shown by the meta-analysis of [14] is close to the 43% reduction in depletion estimated in the BACI experiment by [56].

We used three complementary indicators to assess the impact of beam trawling on the seafloor habitats. The L1 method estimates the biomass proportion of the benthic community with a life span exceeding the average interval between successive trawling events given the observed trawling intensity. As such it is particularly sensitive for changes in trawling intensity in grid cells trawled at low intensity. The L2 method estimates the change in the longevity composition of the benthic community which can be considered to be a proxy for biodiversity. The PD method estimates the decrease in benthic biomass caused by trawling. Since biological activities are scaled to biomass, the biomass method can be considered a proxy for the trawling impact on trophic processes. The PD method additionally allows distinguishing between differences in bottom contact and penetration depth between gear types.

The observed decrease in the L1 and L2 impact indicators is consistent with the observed decrease in trawling footprint and in the PD indicator with the replacement of tickler chains by electrode arrays. The decrease in impact is slightly counteracted by the shift in spatial distribution resulting in a small increase in pulse trawling in coarse sediment (Eunis habitat 5.1). A shift from muddy to coarse sediments will result in a relative increase in benthic impact because coarse sediments have more long lived species than muddy sediments[37].

The response of the L2 indicator to the transition differs between the SFA and the total fishing area (Table 4). The increase in the L2 indicator for the total area can be explained by the interaction of natural disturbance and trawling disturbance events on the benthic community [58, 59]. The empirical relationship between the longevity composition and habitat variables included a significant interaction between bed shear stress and trawling intensity[37]. According to this model, the benthic community in most parts of the southern North Sea is insensitive to beam trawling. Only the benthic communities in areas with low bed shear stress, such as found in the fishing areas north of SFA, are sensitive to trawling. Hence, the increase in beam trawling activities in these areas that are targeting plaice is responsible for the increasing trend in L2. The increase in trawling for plaice is unrelated to the transition to pulse trawling but related to the recovery of the plaice population. Before the collapse of the plaice stock in the early 1990s, these northern grounds were regularly trawled by the Dutch beam trawl fleet [47, 60].

The transition from tickler chain beam trawling to pulse trawling resulted in a substantial reduction in the amount of silt being mobilized. The decrease in sediment mobilization is due to (i) the decrease in towing speed, leading to a reduction in hydrodynamic drag; (ii) the replacement of transverse tickler chains by longitudinal electrode arrays; and (iii) the slight displacement of effort from muddy to coarse sediments. Sediment mobilization has important consequences for the bio-geochemical processes in the sediment – water interface. Sediment mobilization may result in the loss of organic material from the sea bed and a release of nutrients to the overlying water column, while in the water column, the mobilised organic matter may be decomposed by microbial activity [3, 11, 61]. Loss of organic matter due to trawling is of great concern along the continental slope [62], but has also been reported in continental shelf areas [63, 64]. An experimental study of the effect of pulse and tickler chain trawling on biogeochemical processes showed that beam trawling resulted in an immediate decline in benthic community metabolism, with tickler chain trawling exhibiting a stronger effect than pulse trawling [65].

The small reduction of pulse trawling in muddy habitats is in contrast to anecdotal information from the fishing industry suggesting that pulse trawls moved into previously unfished muddy grounds in the southern North Sea[53]. It is possible that the spatial scale used in the present study (1.8 km latitude * 1.1 km longitude at 52°N) is too coarse and may confound habitat differences that occur at smaller scale, such as the pattern of trough’s and ridges which differ in grain size and benthic community [66, 67]. Further analysis at a finer scale is required to resolve this issue.

Our study focussed on the effect of mechanical disturbance on the benthic ecosystem and did not consider the possible effect of electrical pulses. Laboratory studies where benthos was exposed to electrical pulses used in the sole fishery did not find evidence for pulse–induced mortality of a variety of benthic invertebrates [53, 68, 69]. Field and laboratory studies on the effect of pulse trawling and tickler chain trawling on biogeochemical processes only showed biochemical impacts coming from mechanical disturbance but did not find evidence that electrical pulses led to a detectable impact on biogeochemistry (Tiano in [53]). Although studies on the effect of pulse stimuli on marine biota and geochemical processes are still ongoing, the available evidence suggests that the impact of pulse trawls on the benthic ecosystem is due mainly to mechanical disturbance. Hence, replacing mechanical stimulation by tickler chains with electrical stimulation by electrodes will substantially reduce the impact of the beam trawl fishery on the seafloor and the benthic ecosystem when exploiting the Total Allowable Catch of sole.

This study applied, and extended, the mechanistic approach to assessing the physical impact of bottom trawling on the sea floor and the benthic community [33, 70]. This approach integrates quantitative information on the distribution of the trawling activities and the sea floor habitats[4, 5], fishing gear dimensions [35, 40] and the sensitivity of the benthic community[36, 37]. Here we extended the approach by estimating the sediment mobilization due to the hydrodynamic drag in the wake of the gear components, which has important ramifications for the biogeochemical processes [71]. The indicators used to summarise the trawling impacts cover complementary dimensions of the sea floor habitat and benthic ecosystem. The study illustrates the usefulness of the recently developed framework to provide quantitative information on the impact of different fishing gears, which can be used for policy decisions to reduce the impact through gear technological innovations.

## Acknowledgement

This study was in part funded by the EU FP 7 project BENTHIS (grant no. 312088) and the European Maritime and Fisheries Fund (EMFF) through the Netherlands Ministry of Agriculture Nature and Food Quality (LNV) (Grand/Award Number: 1300021172). This article does not necessarily reflect the view of European Commission or the Netherlands Ministries and does not anticipate the Commission/Dutch government’s future policy in this area.

## Author contribution

**Conceptualization**: A.D. Rijnsdorp, J. Depestele, O.R. Eigaard, N.T. Hintzen, A. Ivanovic, F. O’Neill, H. Polet, T. van Kooten

**Data curation**: A.D. Rijnsdorp, J. Depestele, N.T. Hintzen, P. Molenaar

**Formal analysis**: A.D. Rijnsdorp, J. Depestele, N.T. Hintzen, J.J. Poos

**Methodology:** A.D. Rijnsdorp, F. O’Neill, J. Depestele, O.R. Eigaard, N.T. Hintzen, A. Ivanovic,

**Writing – original draft:** A.D. Rijnsdorp

**Writing – review & editing:** A.D. Rijnsdorp, J. Depestele, O.R. Eigaard, N.T. Hintzen, A. Ivanovic, P. Molenaar, F.

**O’Neill**, H. Polet, J.J. Poos, T. van Kooten

